# Male and female variability in response to chronic stress and morphine in C57BL/6J, DBA/2J, and their BXD progeny

**DOI:** 10.1101/2024.02.23.581795

**Authors:** Carole Morel, Lyonna F. Parise, Yentl Van der Zee, Orna Issler, Min Cai, Caleb Browne, Anthony Blando, Katherine Leclair, Sherod Haynes, Robert W. Williams, Megan K. Mulligan, Scott J. Russo, Eric J. Nestler, Ming-Hu Han

**Affiliations:** Department of Pharmacological Sciences, Icahn School of Medicine at Mount Sinai, New York, NY, USA; Friedman Brain Institute, and Center for Affective Neuroscience, Icahn School of Medicine at Mount Sinai, New York, NY, USA; Nash Family Department of Neuroscience, Icahn School of Medicine at Mount Sinai, New York, NY, USA; University of Tennessee Health Science Center.; Department of Mental Health and Public Health, Faculty of Life and Health Sciences, Shenzhen Institute of Advanced Technology, Chinese Academy of Sciences, Shenzhen, Guangdong, PRC

**Author notes:** Equal contribution (Carole Morel, Lyonna F. Parise, Yentl Van der Zee, Scott J. Russo, Eric J. Nestler, Ming-Hu Han).

## Abstract

Drug addiction is a multifactorial syndrome in which genetic predispositions and exposure to environmental stressors constitute major risk factors for the early onset, escalation, and relapse of addictive behaviors. While it is well known that stress plays a key role in drug addiction, the genetic factors that make certain individuals particularly sensitive to stress and thereby more vulnerable to becoming addicted are unknown. In an effort to test a complex set of gene x environment interactions—specifically *gene x chronic stress*—here we leveraged a systems genetics resource: BXD recombinant inbred mice (BXD5, BXD8, BXD14, BXD22, BXD29, and BXD32) and their parental mouse lines, C57BL/6J and DBA/2J. Utilizing the chronic social defeat stress (CSDS) and chronic variable stress (CVS) paradigms, we first showed sexual dimorphism in the behavioral stress response between the mouse strains. Further, we observed an interaction between genetic background and vulnerability to prolonged exposure to non-social stressors. Finally, we found that DBA/2J and C57BL/6J mice pre-exposed to stress displayed differences in morphine sensitivity. Our results support the hypothesis that genetic variation in predisposition to stress responses influences morphine sensitivity and is likely to modulate the development of drug addiction.

## Introduction

Drug addiction is a partly heritable, polygenic disorder determined by a complex interaction between multiple genes and the environment^1^. Familial, twin, and adoption studies have consistently reported that genetic factors contribute to aspects of addiction primarily through interactions with environmental factors and exposure^2,3^. Converging lines of evidence from both clinical and preclinical investigations support the view that environmental stress is a major risk factor for developing drug addiction^4–6^. Indeed, clinical studies strongly suggest that cumulative or prolonged stress is a reliable predictor of drug addiction^5^, and preclinical studies in rodents have confirmed that chronic exposure to stress increases initiation of drug use, induces more robust drug-induced conditioned place preference (CPP), and escalates drug self-administration^5,7,8^.

Although genetic studies have shown addiction to be roughly 50% heritable^9–13^, opioid addiction is accelerating in the US and we are faced with a worldwide problem that accounts for tremendous morbidity and mortality, thereby posing collateral damage to social and political systems^14–17^. Still, the DNA variants that contribute to the behavioral relationship between stress and addiction remain elusive. To this end, the development of expanded families of fully isogenic replicable cohorts^18–20^, in particular, the extended BXD family of recombinant inbred mouse strains generated from a cross between C57BL/6J females and DBA/2J males, have enabled large research efforts in systems genetics and have been used widely to investigate genetic factors underlying complex heritable phenotypes observed in metabolic^21–23^ and psychiatric disorders^24–26^. Each of the ∼120 extant BXD strains are essentially immortal allowing for reliable and reproducible genetic investigations. The BXD family provides a unique system to map the complex set of gene x environment interactions, such as those between genes and stress that are implicated in the vulnerability to drug addiction^26–31^.

In the present study, we investigate such gene-stress interactions in the context of morphine exposure. Utilizing the chronic social defeat stress paradigm (CSDS)^32,33^, we first showed that susceptibility to CSDS varies amongst BXD strains and sexes. To investigate the interaction between genetics and vulnerability to prolonged exposure to non-social stressors, we exposed C57BL/6J, DBA/2J, BXD8, BXD22, and BXD29 male and female mice to the chronic variable stress paradigm (CVS)^34^. We confirmed that behavioral responses to CVS differ among BXD progeny and by sex. Interestingly, we observed that several strains were more vulnerable to non-social chronic stress than chronic social stress. Finally, DBA/2J and C57BL/6J mice pre-exposed to CSDS displayed differences in morphine sensitivity. Characterization of the genetic, neurobiological, social, and environmental factors mediating addiction risk will fundamentally improve our understanding of individual variations in response to drug of abuse and provide highly useful information for developing new individualized prevention and treatment.

## Results

### C57BL/6J, DBA/2J, and BXD mice have distinct sensitivities to chronic social stress

Drug addiction, including morphine-related behaviors, has been linked to several factors such as sex, age, social environment, and stress experience prior to drug exposure^5^. Similar to humans, mice exposed to chronic social stress exhibit a higher propensity to develop addictive behaviors. In our study, we first aimed to determine the effect of genetics and sex on chronic social stress-induced behavioral outcomes. Here we used CSDS, a well-establish mouse model for social stress-induced behavioral alterations^32,35,36^.

We first exposed adult male and female mice from the BXD founders, C57BL/6J and DBA/2J, and the BXD5, BXD8, BXD14, BXD22, BXD29, and BXD32 strains to CSDS (Fig 1a). We assessed their social interaction behavior determined by the social interaction ratio (see detailed information in the methods section). We observed a large range of social interaction ratios across mouse lines and sex (Fig 1b; ANOVA, F_(15,231)_=4.232, p<0.001). Our results showed that DBA/2J and BXD22 male mice are more susceptible to chronic social stress than C57BL/6J mice, as evidenced by their stronger social avoidance behaviors (Fig 1b). Following social interaction, we tested exploratory behaviors in an open-field test (OFT). We observed a large range of stress-induced anxiety-like behaviors across the mouse lines (Fig 1b). In particular, BXD22, BXD29, and BXD8 female mice developed a higher level of anxiety-like behavior than BXD5 and C57BL/6J male mice following CSDS, as evidenced by their decreased time spent in the center of the open field (ANOVA, F_(15,220)_=2.819, p<0.001). We then calculated the distance traveled in the open-field chamber and further observed a range of locomotor activity across mouse lines and sex (ANOVA, F_(15,231)_=21.97, p<0.001).

**Figure 1:**
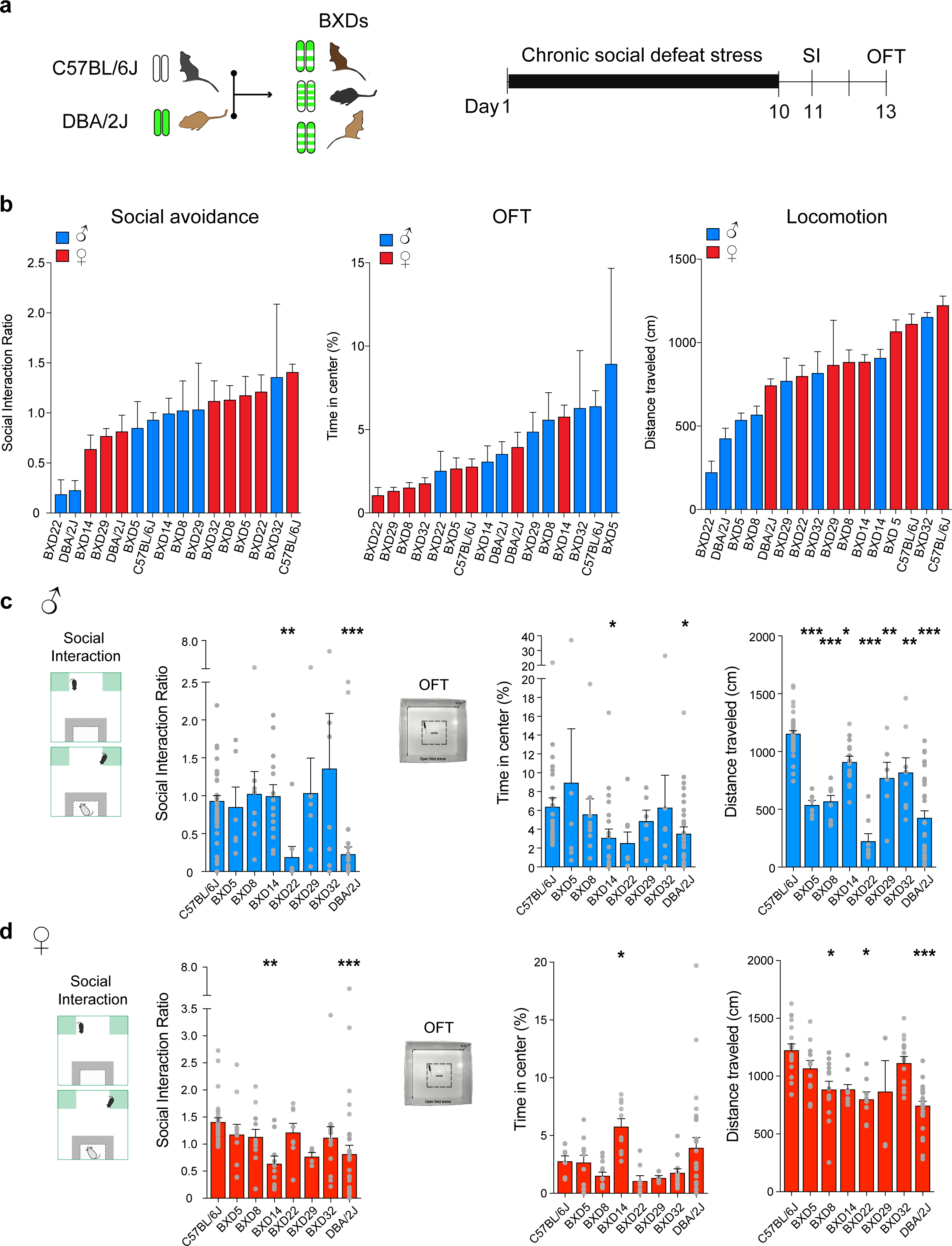
C57BL/6J, DBA/2J, and BXD mice have distinct sensitivity to chronic social defeat stress (CSDS). **(a)** Experimental timeline illustrating the CSDS paradigm followed by the social interaction test (SI) and the open-field test (OFT). **(b)** Behavioral assessment showing the heterogeneous impact of CSDS on SI, OFT, and locomotion by genetic background in male (blue) and female (red) stressed mice. **(c)** CSDS-exposed male mice display distinct SI ratio, percentage of time spent in the center of the OFT, and distance traveled in the OFT in a “no-target” context amongst C57BL/6J (n=40), BXD: 5, 8, 14, 22, 29, and 32 (n=6-14), and DBA/2J (n=35) male mice following CSDS. **(d)** Same as c in CSDS-exposed female mice, C57BL/6J (n=28), BXD: 5, 8, 14, 22, 29, and 32 (n=4-14), and DBA/2J (n=18). Bars represent mean ± SEM. ANOVA test followed by post-hoc comparison to C57BL/6J mice with Bonferroni correction *P<0.05, **P<0.01, and ***P<0.01.

Observing sex differences in response to CSDS amongst the different BXD lines tested, we examined the behavioral responses to CSDS of male mice independently from female mice (Fig 1c,d). We observed that male DBA/2J and BXD22 were more susceptible to CSDS-induced social avoidance than the male founder C57BL/6J line (t-tests, t=2.515, p=0.01; t=3.983, p<0.001). DBA/2J and BXD14 male mice displayed higher CSDS-induced anxiety-like behavior when compared to the male C57BL/6J founder line (t-tests, t=1.987, p=0.04; t=1.886, p=0.04). We also observed that male DBA/2J and BXD lines had lower distance traveled than the male founder C57BL/6J line (t-tests, BXD32 t=3.40, BXD29 t=3.438, BXD14 t=3.096, p=0.02; BXD8 t=6.236, BXD5 t=5.553, BXD22 t=9.442, DBA/2J t=12.16, p<0.001).

Confirming the higher sensitivity of the DBA/2J mice to CSDS-induced social avoidance, female DBA/2J mice had a lower social interaction ratio than C57BL/6J female mice following CSDS (Fig 1d, t=3.434, p<0.001). While BXD14 male mice presented a similar degree of social interaction as male C57BL/6J line, female BXD14 had a lower social interaction ratio than female C57BL/6J mice (t-tests, t=6.727, p<0.001). Additionally, BXD14 female mice displayed higher CSDS-induced anxiety-related behavior when compared to the female founder C57BL/6J line (t-tests, t=2.293, p=0.02). We also observed that female DBA/2J (t-tests, t=3.089, p=0.003), BXD8 (t-tests, t=2.709, p=0.04), and BXD22 (t-tests, t=3.172, p=0.01) lines had lower distances traveled than the female founder C57BL/6J line. Together, these results suggest that CSDS-induced behavioral outcomes are impacted by sex and genetic background.

### C57BL/6J, DBA/2J, and BXD strains have distinct sensitivities to non-social chronic stress

Adverse events are highly heterogeneous, and behavioral and physiological responses to stress vary with the nature of stressors. To capture the impact of genetics and sex on stress-induced behavioral outcomes, we tested behavioral responses of BXD and founder C57BL/6J and DBA/2J mouse lines to the CVS paradigm. We and others have established that exposure to this non-social chronic stress disrupts exploratory behaviors, reward responses, and motivated behaviors ^34,37^.

We first exposed male and female mice from the BXD founder lines C57BL/6J and DBA/2J and the BXD8, BXD22, and BXD29 lines to the CVS paradigm (Fig 2a). We then assessed anxiogenic behaviors in the novelty-suppressed feeding (NSF) task and observed a large range of behaviors across mouse lines and sexes (Fig 2b; ANOVA, F_(9,141)_=9.49, p<0.01). Our results showed that male and female BXD29 mice were more susceptible to CVS-induced NSF than C57BL/6J and DBA/2J mice, as evidenced by their elevated latency to feed (Fig 2b). We then tested the extent of CVS-induced anhedonia by measuring the preference for sucrose solution over water. We observed a stronger impact of CVS on BXD22 and BXD8 sucrose preference (ANOVA, F_(9,143)_=2.73, p=0.006). Following these measurements, we tested exploratory behavior in an elevated plus maze (EPM) and observed a large range of stress-induced anxiety-like behaviors across mouse lines and sexes (Fig 2b; ANOVA, F_(9,143)_=6.468, p<0.001). In particular, BXD22 and BXD29 mice developed a higher level of anxiety than C57BL/6J and DBA/2J mice after CVS, as evidenced by lower time spent in the open arms of the EPM. Further confirming the variable sensitivity to stressful stimuli, we observed a large range of locomotor behaviors across mouse lines and sex (Fig 2b; ANOVA, F_(9,143)_=19.36, p<0.001).

**Figure 2:**
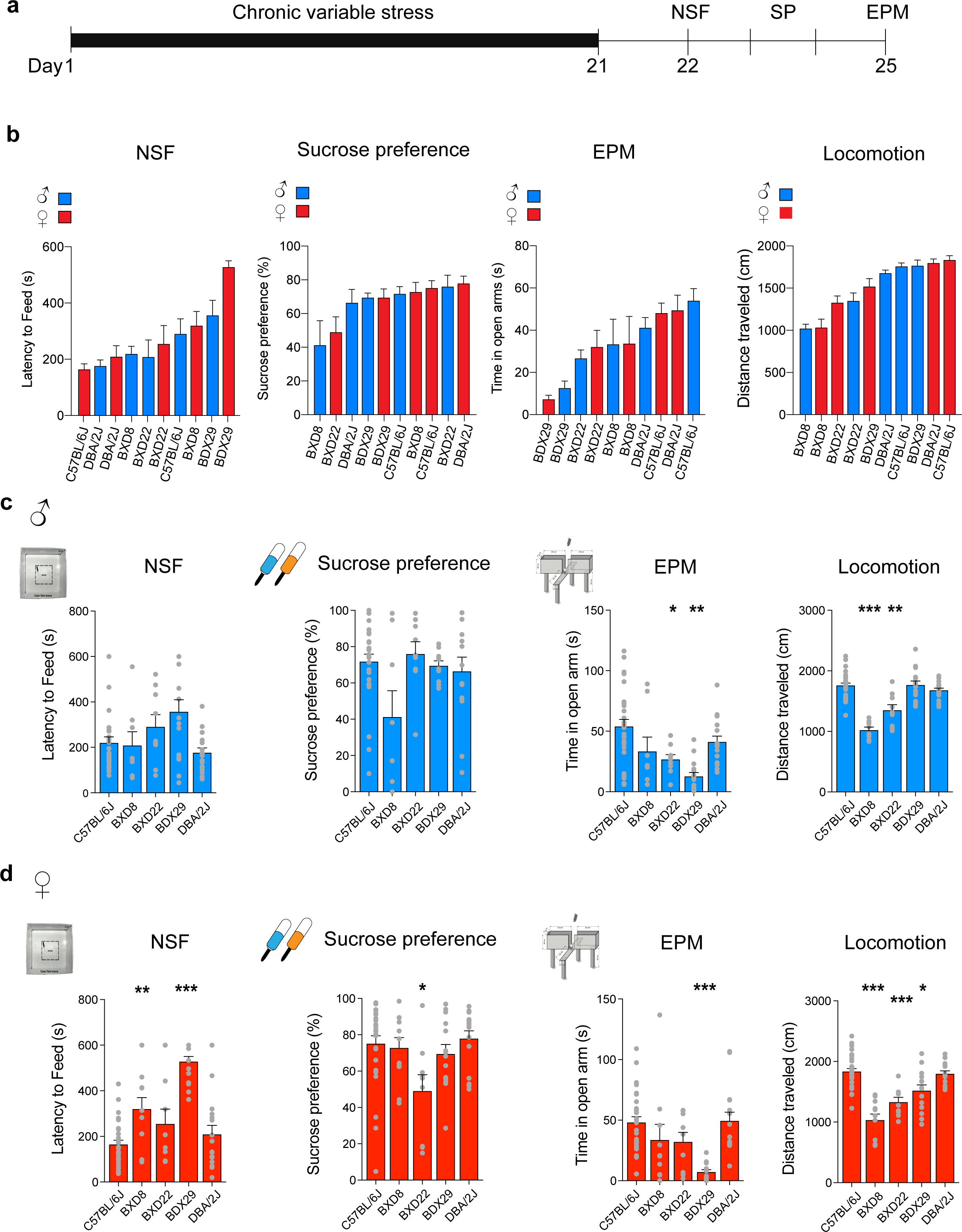
C57BL/6J, DBA/2J, and BXD mice have distinct sensitivity to non-social chronic stress. **(a)** Experimental timeline illustrating the chronic variable stress (CVS) paradigm followed by the novelty suppress feeding test (NSF), the sucrose preference test (SP), the elevated plus-maze test (EPM), and locomotion in an open field. **(b)** Behavioral assessment showing the heterogeneous impact of CVS on NSF, SP, EPM, and locomotion by genetic background. Bars represent mean ± SEM for n=6-40 mice. **(c)** CVS-exposed male mice display distinct latency to feed during the NSF task, SP ratio (%), time spent as percentage in open arms of the EPM, and distance traveled in an open field in the “no-target” context amongst C57BL/6J (n=27), BXD: 8, 22, and 29 (n=9-11) and DBA/2J (n=13) male mice. **(d)** CVS-exposed female mice display distinct latency to feed times during the NSF task, SP ratio (%), time spent in EPM open arms, and distance traveled in an open field in “no-target” context amongst C57BL/6J (n=27), BXD: 8, 22, and 29 (n=8-15) and DBA/2J (n=16). Bars represent mean ± SEM. ANOVA test followed by posthoc comparison to C57BL/6J mice with Bonferroni correction *P<0.05, **P<0.01, and ***P<0.01.

We did not observe significant differences in NSF and sucrose preference ratio between males following CVS exposure (Fig 2c). Yet, BXD29 and BXD22 males had higher symptom levels when compared to the male founder C57BL/6J line (t-tests, t=2.293, p=0.04; t=2.293, p=0.002; Fig 2c). We also observed that male BXD8 (t-tests, t=8.477 p<0.001) and BXD22 (t-tests, t=4.92 p<0.001) lines had lower distances traveled than the male founder C57BL/6J line (Fig 2c).

Confirming the heterogeneous stress-induced behavioral responses between sex and strain, we observed that female BXD29 and BXD8 mice displayed a higher latency to feed when compared to C57BL/6J female mice (Fig. 2d, t=8.640, p<0.001, t=3.211, p=0.006). While sucrose preference in BXD22 males was similar to C57BL/6J males, BXD22 females had a lower sucrose preference ratio relative to C57BL/6J females (t-tests, t=3.224, p=0.008). In addition to high NSF, BXD29 females displayed higher CVS-induced anxiety-like levels when compared to C57BL/6J females (t-tests, t=4.807, p<0.001). We also observed that BXD8 (t-tests, t=8.141, p<0.001), BXD22 (t-tests, t=4.766, p<0.001), and BXD29 (t-tests, t=3.484, p=0.002) females had lower distances traveled relative to C57BL/6J females. Together these results confirm that sex and strain have a relevant influence on chronic social and non-social stress behavioral responses.

### Distinct impact of chronic stress on morphine-induced motor effects between C57BL/6J and DBA/2J mouse lines

Variability in response to psychoactive drugs in mice is known to depend in part on genetic differences^38–41^. To test if genetics and sex impact the chronic stress effects on morphine sensitivity, we measured morphine-induced motor activity in stressed male and female C57BL/6J and DBA/2J mice.

Following the CSDS or CVS paradigm, male and female C57BL/6J and DBA/2J mice were injected with 7.5 mg/kg of morphine (*i.p.*; Fig 3a). This dose was selected because it consistently and reproducibly produces a sub-maximal degree of behavioral responses in C57BL/6J mice (males and females), making it possible to detect manipulations that either increase or decrease sensitivity to the place conditioning effects of the drug. Fifteen minutes after the morphine injection, mice were placed in a large cage allowing for tracking of their locomotor activity. We observed that male and female C57BL/6J mice exposed to CSDS had higher locomotor activity when compared to DBA/2J mice (Fig 3b; t-tests, t=6.452, p<0.001; t-tests, t=6.960, p<0.001).

**Figure 3:**
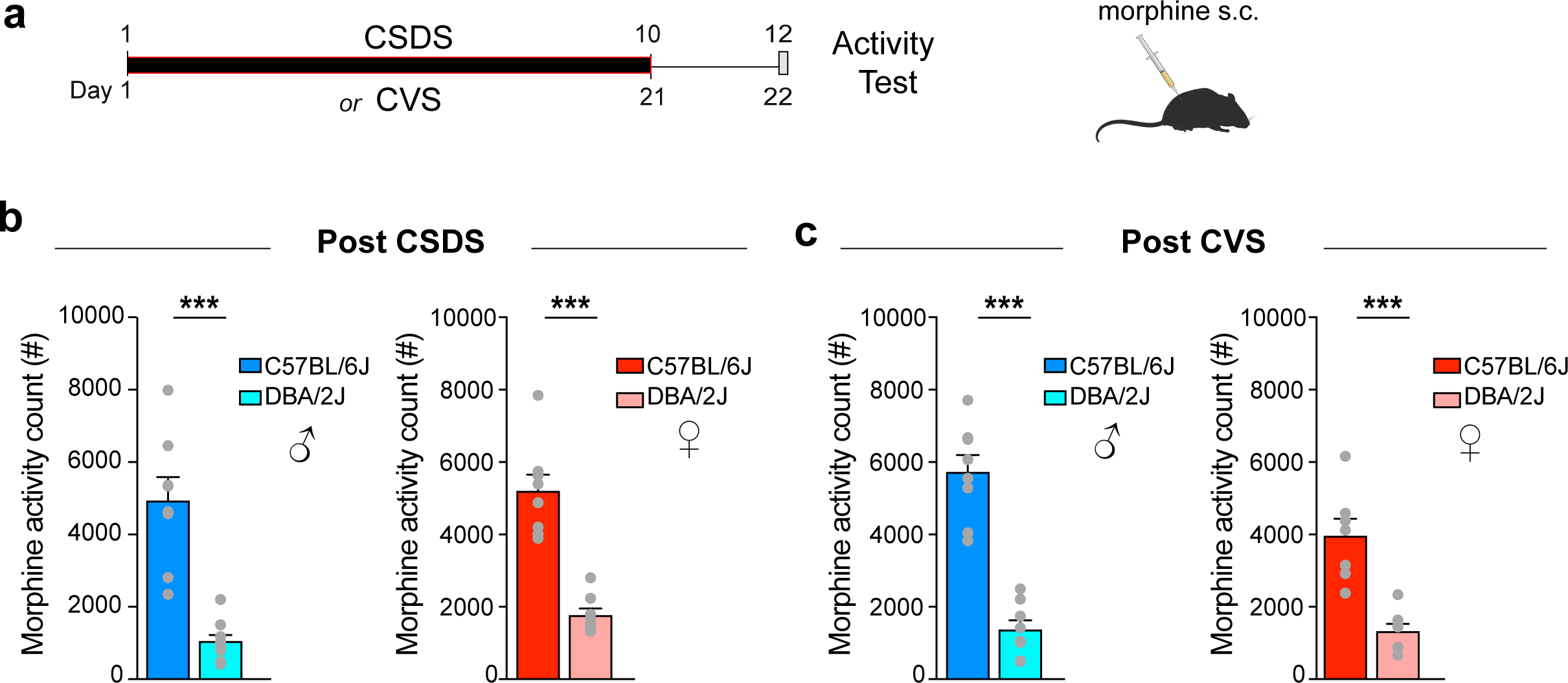
Parental C57BL/6J and DBA/2J mice have distinct locomotor responses to acute morphine after chronic stress exposure. **(a)** Experimental timeline illustrating the two procedures performed, *i.e.* CSDS or CVS, followed by locomotor activity monitoring for 60 min in a locomotor apparatus 15 min after the animals were given an *i.p.* injection of 7.5 mg/kg morphine. **(b)** Locomotor activity counts in C57BL/6J and DBA/2J male (left) and female (right) mice (n=8) exposed to CSDS. **(c)** Locomotor activity counts in C57BL/6J and DBA/2J male (left) and female (right) mice (n=8) exposed to CVS. Bars represent mean ± SEM. Kruskal-Wallis test followed by post-hoc comparison to C57BL/6J mice with Bonferroni correction and non-paired *t*-test, *P<0.05, **P<0.01, and ***P<0.01.

Recapitulating these data, male and female C57BL/6J mice exposed to CVS had higher locomotor activity when compared to DBA/2J mice (Fig 3c; t-tests, t=8.080, p<0.001; t-tests, t=5.301, p<0.001). Together, our results confirmed that the interaction between chronic stress and behavioral responses to morphine vary between genetic backgrounds.

### Sex-specific impact of chronic stress on morphine’s rewarding effects in C57BL/6J and DBA/2J mouse lines

To test the hypothesis that chronic stress divergently impacts the rewarding properties of morphine between C57BL/6J and DBA/2J male and female mice, we employed the CPP paradigm (Fig 4a). Following CSDS or CVS, male and female C57BL/6J and DBA/2J mice were conditioned with once-daily alternating injections of saline and morphine (7.5 mg/kg, *s.c.*) over a period of 2 days. Side preference was assessed one day prior to morphine conditioning (pretest), and an unbiased CPP approach was utilized. One day after conditioning, the preference for the morphine-paired compartment was assessed.

**Figure 4:**
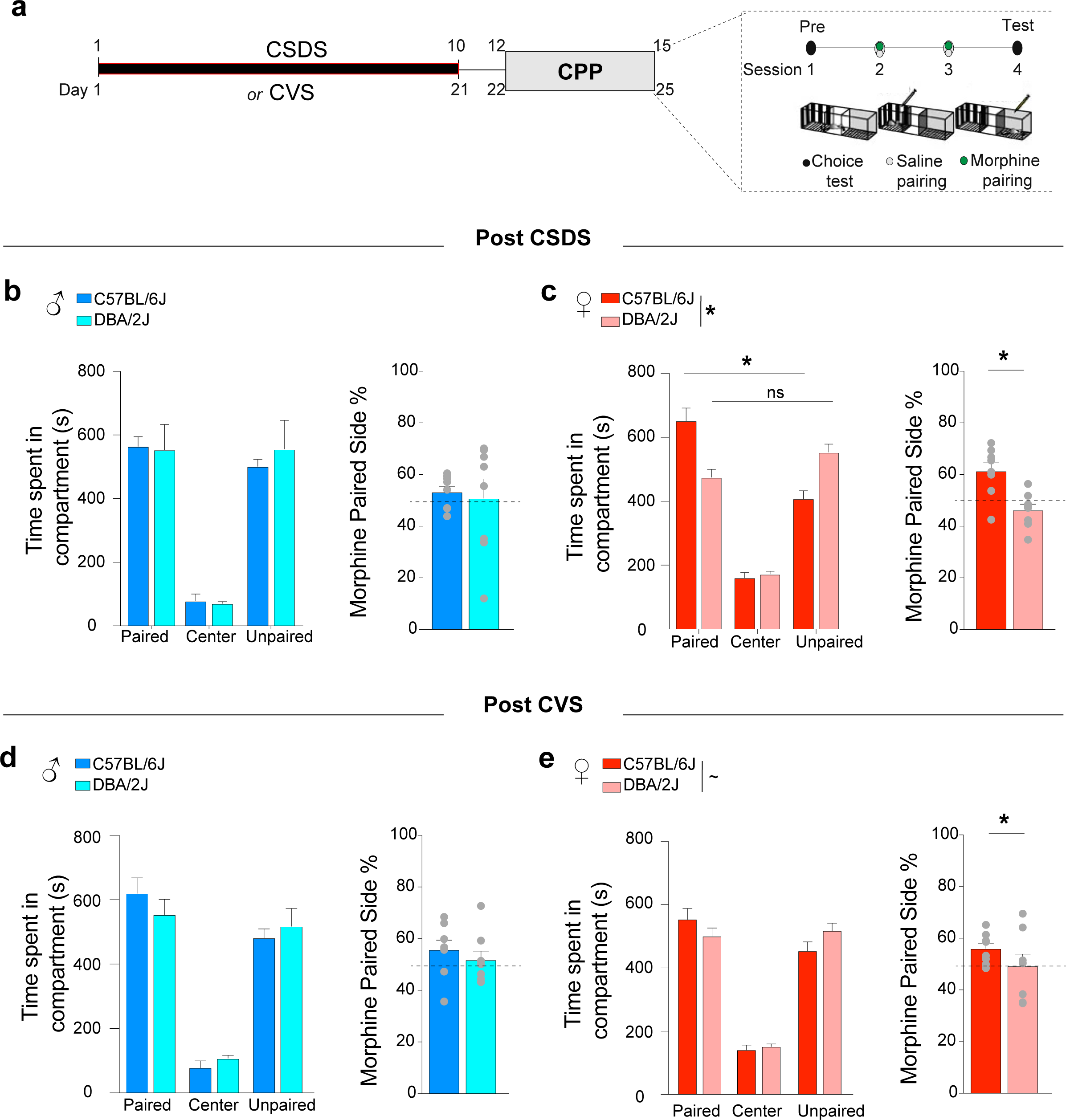
Parental C57BL/6J and DBA/2J mice have distinct place conditioning responses to morphine after chronic stress exposure. **(a)** Experimental timeline illustrating the two procedures performed, *i.e.* CSDS or CVS, prior to performing morphine conditioned place preference (CPP). Morphine CPP was performed using an unbiased approach: one compartment was paired with saline (0.3 mL, *s.c.*) and the other paired with morphine (7.5 mg/kg, *s.c.*). Sessions were 20 min long. **(b)** Time spent in each compartment of the three-chamber apparatus after paired-conditioning with morphine (vs. saline) in CSDS-exposed C57BL/6J and DBA/2J male mice (n=8) and respective percentage in the morphine-paired chamber. **(c)** Same as b, in CSDS-exposed C57BL/6J and DBA/2J female mice (n=8). **(d)** Time spent in each compartment of the three-chamber apparatus after paired-conditioning with morphine (vs. saline) in CVS-exposed C57BL/6J and DBA/2J male mice (n=8) and respective percentage in the morphine-paired chamber. **(e)** Same as b, in CVS-exposed C57BL/6J and DBA/2J female mice (n=8). Bars represent mean ± SEM. Kruskal-Wallis test followed by post-hoc comparison to C57BL/6J mice with Bonferroni correction and non-paired t-test, *P<0.05, **P<0.01, and ***P<0.01.

We observed that CPP scores and preferences did not show significant differences between male CSDS-exposed C57BL/6J and DBA/2J mice (Fig 4b; TWO-WAY ANOVA, F_(2,42)_=0.25, p=0.77; Mann Whitney, U=28, p=0.72). TWO-WAY ANOVA comparison showed that, while CSDS-exposed female C57BL/6J mice established morphine CPP, CSDS-exposed female DBA/2J mice did not establish a preference for the morphine-paired compartment (Fig 4b; TWO-WAY ANOVA, F_(2.42)_=20.14, p<0.0001; paired-side t=4.886 p<0.001, unpaired-side t=4.047 p=0.007). We further observed significant differences in preference for the morphine-paired compartment between CSDS female C57BL/6J and DBA/2J mice (Fig 4c; t-test, t=3.676, p=0.002). Recapitulating these results, CVS-exposed female C57BL/6J and DBA/2J mice showed trending differing CPP profiles (Fig 4d; TWO-WAY ANOVA, F_(2,42)_=3.19, p=0.059), whereas CVS-exposed male C57BL/6J and DBA/2J mice did not (Fig 4e; TWO-WAY ANOVA, F_(2,42)_=1.079, p=0.34). Additionally, CVS-exposed male C57BL/6J and DBA/2J mice did not show a significant difference in preference for the morphine-paired side (Fig 4d; t-test, t=0.791, p=0.44), and CVS-exposed female C57BL/6J mice established CPP, while DBA/2J females did not (Fig 4e; t-test, t=2.17, p=0.045). Our results establish a strain- and sex-specific impact of chronic stress on the rewarding properties of morphine in C57BL/6J and DBA/2J mouse lines.

## Discussion

While stress is a well-known risk factor in the development of drug addiction, the genetic factors that make certain individuals particularly susceptible or resilient to stress and, thereby, more or less vulnerable to becoming addicted, remain elusive. In this study, we tested if genetic components map onto the behavioral and physiological mechanisms underlying 1) the vulnerability to chronic stress exposure, and 2) the ability of prior stress experience to influence drug responses. The BXD family of recombinant inbred strains derived from crossing two inbred parental strains—C57BL/6J and DBA/2J mice—have been extensively used for almost 50 years in fields such as neuropharmacology^26^, immunology^42^, and addiction^18,25,39^ to answer important biological questions. Combining the use of the BXD mouse lines and their founder lines with models of chronic stress exposure and morphine sensitivity, we first showed that CSDS-induced behavioral outcomes are impacted by sex and strain. Further, we established that vulnerability to chronic non-social stress (*i.e.*, CVS) also varies depending on sex and genetics. We revealed that BXD22 male and female mice are more susceptible to chronic social stress than C57BL/6J mice, evidenced by stronger social avoidance and anxiety-like behaviors. We observed sexual dimorphism in responses to CSDS amongst the BXD5, BXD8, BXD14, BXD29, and BXD32 lines. To investigate the interaction between genetics and vulnerability to prolonged exposure to non-social stressors, we exposed C57BL/6J, DBA/2J, BXD8, BXD22, and BXD29 male and female mice to CVS and observed that DBA/2J female mice are more sensitive to CVS when compared to C57BL/6J female mice (*i.e.*, a heightened decrease in sucrose preference). Confirming the stress vulnerability of BXD22 mice observed after CSDS, CVS-exposed male and female BXD22 mice displayed higher levels of anxiety-like measures than C57BL/6J mice. Interestingly, while BXD29 mice behaved like C57BL/6J after CSDS, both BXD29 female and male mice developed a higher anxiety profile following CVS when compared to C57BL/6J mice. Finally, we identified that DBA/2J and C57BL/6J mice pre-exposed to CSDS displayed differences in morphine sensitivity.

Multiple studies, including our own, have established that CSDS decreases social interaction behaviors, even to the point of avoidance, and reduces exploratory behaviors. We observed that DBA/2J and C57BL/6J mice have distinct behavioral responses following CSDS exposure, with DBA/2J mice exhibiting stronger social avoidance and lower exploratory behaviors when compared to C57BL/6J mice. Interestingly, following CVS exposure, DBA/2J and C57BL/6J mice exhibited similar behavioral responses. Conversely, while BXD29 mice behaved like C57BL/6J mice after CSDS, the BXD29 mice developed a heightened anxiety profile following CVS compared to C57BL/6J mice. These results suggest distinct genetic contributions in the mechanisms underlying these two stress-induced behavioral outcomes. In line with these observations, several studies have reported distinct and even opposite modifications of neural circuits involved in social and non-social stress exposure, including within the dopamine system^32,43,44^. In particular, it has been shown that CSDS induces a hyper-dopaminergic state in mice expressing social avoidance and decreased sucrose preference. Oppositely, chronic non-social stress reduces the activity of midbrain dopaminergic neurons in mice exhibiting depressive-like behaviors such as a lower sucrose preference ratio. Together, these results emphasize that opposite neuronal modifications, supported by distinct genetic mechanisms, may converge to similar pathological behavioral responses to distinct types of stressors.

In line with evidence in humans^34,45,46^, we observed sexual dimorphism across the parental and BXD mouse lines in behavioral outcomes after chronic stress exposure. For example, we observed that, while male BXD22 mice developed stronger social avoidance behaviors compared to male C57BL/6J mice, female BXD22 mice did not and maintained social behaviors similar to those of female C57BL/6J mice exposed to CSDS. Notably, we observed that CSDS-exposed female mice had lower exploratory behaviors in the center of the open-field when compared to their male counterparts. Interestingly, as a result of prolonged exposure to non-social stressors, we observed similar behavioral alterations between male and female mice within the mouse lines. Both male and female BXD29 mice exhibited the strongest stress-induced behavioral deficits compared to C57BL/6J mice. These results indicate stress-specific sex differences in the mechanisms underlying behavioral responses to these two types of stress.

Similar to the behavioral changes after chronic stress exposure, sex is a key factor that may play a role in the development of addiction. Women tend to progress more rapidly than men through the stages of substance use disorder despite initiating drug use at a later age^47^. Women are more likely to report misusing prescription opioids to cope with negative affect, while also showing the characteristic rapid progression from first drug experience to developing substance use disorder^48–50^. To allow for the detection of subtle but robust sex differences in behavioral responses to morphine, we utilized a threshold dose of morphine (7.5 mg/kg), below the dose known to induce CPP or somatic withdrawal in stress-naïve mice^51^. We observed a robust strain effect on morphine-induced locomotor activity between the C57BL/6J and DBA/2J mice. Additionally, female C57BL/6J mice showed greater CPP when compared to female DBA/2J mice, while male DBA/2J and C57BL/6J mice did not establish morphine CPP at the dose used. We found that strain and sex contributed to stress-induced sensitivity to morphine reward, independently from the nature of the chronic stressor, i.e., social or non-social. Further genetic analyses seeking to identify the specific genetic loci involved in the impact of stress on morphine sensitivity will help identify novel mechanisms in gene x environment interactions in neuropsychiatric disorders.

To conclude, our results support the hypothesis that genetic variations in predisposition to stress responses influence sensitivity to morphine and, as a result, presumably modulate the risk of addiction. Our previous studies performed in isogenic mouse lines have allowed the identification of physiological processes and epigenetic mechanisms contributing to stress-susceptibility and drug responses on a single genetic background^52–55^. In parallel, important genetic mapping research efforts using recombinant inbred mouse models have identified the high degree of genetic homology between humans and mice^56^. This homology has allowed for impactful cross-species studies that will define the genes interacting in the regulation of behavioral stress responses and addictive behaviors. Characterization of the genetic, neurobiological, social, and environmental factors that mediate addiction risk will fundamentally improve our understanding of individual variations in responses to drugs of abuse and provide highly useful information for the development of new treatment strategies and, eventually, prevention measures.

## Material and Methods

### Mice

C57BL/6J, DBA2J, and BXD female and male mice (7–12 weeks old; Jackson Laboratory, Bar Harbor) were used for all experiments and habituated to the animal facility for one week before experimental manipulations. Mice were housed in groups of five at a constant ambient temperature (24 +/-1℃) under a 12-hour light/dark cycle (lights on from 7:00 A.M.) with ad libitum access to water and food, except when otherwise specified for behavioral testing. Experiments were conducted in accordance with the guidelines of the Institutional Animal Care and Use Committee (IACUC) at Mount Sinai. Female estrous cycle was monitored by microscopy of vaginal smears at the end of all experiments and confirmed that the large majority of mice were in diestrus.

### Chronic social defeat stress (CSDS) paradigm

Pre-screened CD-1 retired male breeder mice were singled-housed in one compartment of a large mouse cage partitioned by a perforated Plexiglass divider vertically placed through the cage, which has a metal top to hold food and water on both sides. This is used as the resident cage and is the permanent home for the CD-1 mouse. The CSDS paradigm begins when an experimental mouse is introduced into the CD-1 home cage and the ensuing fighting is allowed to occur for 10 min. Following the interaction, mice are individually confined to one compartment for the rest of the day, allowing for continued sensory exposure. This is repeated for 10 days, and the effect of the 10-day defeat is measured on day 11 through a social interaction test^32^. Similar to the male CSDS paradigm, female mice are placed, as an intruder, in the home cage of an Esr1-Cre mouse. The Esr1-Cre mouse is injected with CNO, an exogenous drug targeting the DREADD expressed in the ventral medial lateral hypothalamus (vl), to induce aggressive behavior toward the female experimental mice. Following 5 min of physical contact, the female mice are placed back into their home-cage, where they are grouped housed. Similar to the male CSDS paradigm, this schedule is repeated for 10 days^33,57^.

### Social interaction test (SI test)

Mice were placed in an OFT box containing a wire mesh cage on one side under red-light conditions (<15 lux)^32^. The open field arena and wire-mesh enclosures were thoroughly cleaned between mice with an odorless 5% ethanol solution. The CSDS-exposed mice were individually placed in the open field box and allowed to explore for 2.5 min (without a social target present, known as the “no target” phase”). The experimental mouse was then removed, the open field was cleaned, and an unfamiliar aggressor (CD-1/Esr1-Cre) mouse was placed into the mesh cage (known as the “target phase”). The experimental mouse was then allowed to explore for another 2.5 min. Time spent interacting with the social target and locomotion were measured using an automated video tracking system (Ethovision). Experimental mice were then placed back into their home cage (single-housed). The social interaction ratio was calculated as [time in social interaction zone with “target”]/[time in social interaction zone with “no target”], as described previously^32,58^. The distance traveled was analyzed during the “no target” phase to avoid potential biases due to novel social target being present.

### Chronic variable stress (CVS)

CVS consists of three different stressors over 30 days^34,59^. Environmental stressors are presented on an alternating schedule to prevent habituation. Stressors are administered in the following order: 100 random mild foot shocks at 0.45 mA for one hour (10 mice to a chamber), a tail suspension stress for one hour, and restraint stress (placed inside a 50mL falcon tube) for one hour. The three stressors are then repeated in the same cycle for the remainder of the stress protocol.

### Open field test (OFT)

Mice were placed in the open field arena (44 x 44 cm) for 5 min to compare the distance traveled and time spent in the peripheral zone compared to the center zone (10 x 10 cm). Testing conditions occurred under red-light conditions (<10 lux) in a room isolated from external sound sources. The apparatus was thoroughly cleaned between mice with an odorless 5% ethanol solution. The mouse’s time spent in specific open field areas— was video-tracked and scored with Ethovision software^32,35^.

### Elevated plus maze test (EPM)

The EPM was designed in black Plexiglass (L/W/D: 70/5/20cm) and fitted with white surfaces to provide contrast. Testing conditions occurred under red-light conditions (<10 lux) in a room isolated from external sound sources. The EPM apparatus was thoroughly cleaned between mice with an odorless 5% ethanol solution. Mice were positioned in the center of the maze, and behavior was video-tracked for 5 min^32,35^. Time in EPM compartments was measured using a video-tracking system (Ethovision) set to focus on the mouse center-point at the commencement of each trial.

### Sucrose preference (SP)

Mice were habituated to having access to two water bottles (50-ml tubes with fitted ball-point sipper tubes) for one day, and were then given free access to both water and a 1% sucrose solution, for two consecutive days^32,35,53^. Bottles were weighed daily and interchanged (left to right, right to left) to avoid biases from a potential side preference. Sucrose preference scores were calculated as ([sucrose solution consumed] / [sucrose + water solutions consumed]) x 100.

### Novelty suppressed feeding (NSF)

The NSF test elicits conflicting motivations in mice: the drive to eat vs. the fear of moving to the center of a novel, open arena^45^. Mice were food restricted for 24 hours prior to testing^34^. Under red-light conditions, mice were placed in the corner of a clear plastic testing chamber (50 x 50 x 20 cm) which was lined with corncob bedding. A single food pellet was present in the center of the box. The latency to begin consumption in a 6-min test was measured.

### Morphine conditioned place preference (CPP)

Morphine CPP was conducted as previously described^55^. We used an unbiased, three-compartment apparatus. First day 0, mice were allowed to freely explore the entire apparatus for 30 min to obtain baseline preference to any of the three compartments; we know that mice show no inherent group biases for a given chamber across multiple strains^52,54^. Mice were then given conditioning trials (two per day) on two consecutive days. For both trials, mice received saline (0.3 mL, *s.c.*) and were confined to one of the compartments of the apparatus.

After 4 hr, mice received morphine (7.5 mg/kg in 0.3 mL, *s.c.*) and were confined to the opposite compartment. The drug-paired chamber is randomized across mice. On test day (day 4), mice are again allowed to freely explore the entire apparatus for 30 min. CPP scores are calculated as the time spent in the morphine-paired chamber minus the time spent in the saline-paired chamber. The time spent in each compartment was determined using an automated system and CPP scores were calculated as the percent of time spent in the morphine-paired compartment.

### Locomotor sensitization

Locomotor activity was individually monitored in specialized locomotor chambers with no bedding, 15 min after the animals were given an *i.p.* injection of 7.5 mg/kg morphine. Locomotor activity was monitored for 60 min.

### Data analyses and statistics

All behaviors were monitored and scored using automated and unbiased Ethovision software^32,46^. Experimenters analyzing the dataset were blinded to the experimental conditions. The statistical analyses were performed using Graphpad Prism (version 8, La Jolla, CA, USA) and R (version 3.3.3) software. The normality of the distributions was assessed using Kolmogorov-Smirnov tests. The statistical analyses were performed considering the sample size, normality, and homoscedasticity of the distributions. The data fitting assumptions of the general linear model were subjected to two-sided Student’s *t*-tests, or multiple comparisons using a one-way, two-way, or repeated measures (RM) ANOVA followed by post hoc two-sided t-tests with Bonferroni correction for multiple comparisons (t-test, p-values). Non-parametric Kruskal-Wallis and Mann-Whitney two-sided statistical analyses were performed for datasets that did not follow a normal distribution or homoscedasticity. The statistical significance threshold was set at 0.05.

## Data availability

All data reported by this study are included within the manuscript’s figures or provided in the supplementary information section and Source Data files. Source Data are provided with this paper. Any additional data and information are available upon reasonable request.

## Acknowledgments

This work was supported by National Institute of Mental Health grants R01MH051399 (EJN) and R01MH120514 (CM, SJR) and by National Institute on Drug Abuse grant P01DA047233 (EJN), National Key R&D Program of China 2021ZD0202900 & 2021ZD0202902 (MHH), Research Fund for International Senior Scientists T2250710685 (MHH), Shenzhen Natural Science Foundation J20220127 (MHH) Shenzhen Medical Research Fund SMRF B2303012 (MHH), Shenzhen Key Laboratory of Precision Diagnosis and Treatment of Depression ZDSYS20220606100606014 (MHH), Science and Technology Research and Development Foundation of Shenzhen (High-level Talent Innovation and Entrepreneurship Plan of Shenzhen Team Funding) KQTD20221101093608028 (MHH). This study was also supported by a NARSAD Young Investigator Grant from the Brain & Behavior Research Foundation (CM and LFP 31194), Leon Levy Foundation (LFP) and the Hope for Depression Research Foundation (EJN).

## Authors’ contributions

C.M., L.P., Y.V.Z., O.I., M.C., K.C., A.B., performed the behavioral assessments and with the assistance of C.B., C.M., L.P. Y.V.Z., M.C., O.I., analyzed the results. C.M., L.P., S.J.R., E.J.N., and M.H.H. designed the experiments with the assistance of R.W.W. M.K.M., C.M., L.P., O.I., S.J.R., E.J.N., and M.H.H. interpreted the results and wrote the paper, which was edited by all authors.

## Corresponding authors

Correspondence and requests for materials should be addressed to Ming-Hu Han.

